# Fagalean phylogeny in a nutshell: Chronicling the diversification history of Fagales

**DOI:** 10.1101/2023.03.06.531381

**Authors:** Carolina M. Siniscalchi, Julian Correa-Narvaez, Heather R. Kates, Douglas E. Soltis, Pamela S. Soltis, Robert P. Guralnick, Steven R. Manchester, Ryan A. Folk

**Affiliations:** Department of Biological Sciences, Mississippi State University, Mississippi State, MS, United States; Florida Museum of Natural History, University of Florida, Gainesville, FL, United States; Department of Biology, University of Florida, Gainesville, FL, United States; Genetics Institute, University of Florida, Gainesville, FL, United States; Biodiversity Institute, University of Florida, Gainesville, FL, United States

**Keywords:** biogeography, Cretaceous, fossilized birth-death process, total evidence dating

## Abstract

Reconstructing the biogeographical history and timing of the diversification of temperate forests is essential for understanding their history and resolving uncertainties about how flowering plants emerged from their deep tropical origins to dominate in today’s freezing terrestrial environments. The angiosperm order Fagales, comprising iconic components of temperate forests worldwide with an extensive fossil record, are an excellent plant system in which to apply a fossil-aware paradigm, such as the fossilized birth-death (FBD) process, for investigating the macroevolution of temperate forest biomes. Here, we improve upon previous efforts to resolve phylogeny and incorporate fossils in Fagales using low-copy nuclear loci and an expanded morphological matrix to reevaluate the Fagales fossil record and: (1) infer the phylogenetic relationships and the time of origin of the clade using the FBD model as implemented in RevBayes, (2) provide a framework for evaluating the climatic and biogeographic history of Fagales, and (3) investigate how the inclusion of fossils via the FBD method influences ancestral reconstruction and diversification estimation. The phylogenetic relationships we recovered are conventional except for the position of Nothofagaceae, while our inferred ages support older timelines than previously proposed, with a mid-Cretaceous date for the most recent common ancestor (MRCA) of the order. Biogeographical analysis shows an origin of Fagales consistent with an ancestral circumboreal temperate distribution corroborated by ancestral niche reconstructions. While distributions today largely reflect the general conservatism of temperate forests, we identified two episodes of high diversification, one at the mid-Cretaceous origin of the clade and the other continuing from the Miocene to the present. Removing fossil taxa from the tree reveals a different story, shifting the origin of extant families from North America to East Asia, reflecting refugial distributions in this biodiversity “museum” and implying a general bias towards low extinction areas in biogeographic reconstruction. Likewise, without fossil data, diversification estimates were higher and unable to detect an early diversification burst. Based on our analyses, we close with recommendations regarding the interpretation of estimates of diversification and ancestral state reconstruction using phylogenetic trees with only extant species as tips.

---

Reconstructing the biogeographical history and timing of the diversification of temperate plants is essential for understanding the origins of today’s colder, more extreme environments in the face of a plant evolutionary legacy largely characterized by tropical origins (Folk et al. 2020). Basic uncertainties remain about the age of temperate and cold ecological niches and where these environments may have existed on Earth. A revised timeline of cold tolerance implies very different scenarios for how plants invaded these environments, with an older origin being consistent with the compelling classic hypothesis of co-option from ancient strategies for surviving in seasonal tropical environments (Zanne et al. 2014; Folk et al. 2020). Alternatively, a younger origin of survival in subfreezing conditions would suggest a greater degree of evolutionary novelty (Spriggs et al. 2015; Edwards et al. 2017) but would also raise questions about the origin of groups that today specialize in temperate habitats but originated before such cold habitats were widespread (Folk et al. 2021b). Ancient groups occurring in modern temperate environments are therefore of key interest for understanding the broader question of how plants cope with extremity and global change.

Despite intense recent interest in more fossil-aware macroevolutionary approaches, there remain few studies fully incorporating information from the fossil record to infer the timeline of diversification of taxa. However, this information is critical for addressing broader macroevolutionary questions such as the origin of temperate floras. Calibration-based divergence time estimation remains standard practice even in plant lineages with a rich fossil record that would enable total-evidence approaches (Ronquist et al. 2012). Most studies investigating plant diversification, biogeography, and macroevolution make incomplete use of fossil information, given not only the known limitations of extant-only phylogenies (Rabosky 2010; Louca and Pennell 2020; Helmstetter et al. 2021), but more importantly the rich morphological and ecological information that fossils can contribute to ancestral reconstruction and other phylogenetic applications. Total evidence dating is characterized by treating fossils as sampled taxa in the phylogeny and simultaneously inferring topology and divergence times, fully incorporating the phylogenetic evidence that fossils possess (Ronquist et al. 2012, Heath et al. 2014). This approach permits estimating uncertainty as well as a straightforward path to extracting the ecological and morphological information from fossils by treating them no differently than extant taxa. Such analyses, integrating living and extinct taxa, remain more common in animals with relatively few case studies in plants demonstrating the use of total-evidence dating approaches for understanding plant evolution (Hardy 2006; Wood et al. 2013; Holland et al. 2020; May et al. 2021; Wisniewski et al. 2022).

Biogeographical and diversification analyses benefit greatly from total-evidence dating because inclusion of fossil evidence allows more direct inference of parameters while overcoming the possibility of bias through extant-only timetrees. Models that indirectly take into account geological (rather than fossil) information have been previously proposed and may be effective when used for taxa restricted to areas where the geological history is well known, such as the Hawaiian islands (e.g., Landis et al. 2018). Given the limited utility of this type of information for many continental radiations, ancestral area reconstructions are more commonly based only on current distributions with fossil information incorporated as calibration points for chronograms. Consequently, important parts of the biogeographical history of plant taxa are effectively excluded from the synthesis even when appropriate fossil information is available. Certainly there are cases where fossil geographical distributions accord well with the results of extant-only biogeographic modeling (Landis et al. 2021), but one direct effect of excluding fossils from biogeographical reconstruction is that geographical regions that suffered high rates of past extinctions are totally excluded from the reconstruction, resulting in the potential for bias towards geographical regions of low extinction (Hardy 2006; Wood et al. 2013; Holland et al. 2020; Wisniewski et al. 2022). Such analyses often recover high statistical support, but ultimately have underestimated uncertainty and bias in their reconstructions.

Sampling extinct taxa considered phylogenetically proximal to nodes of interest is likely to greatly improve ancestral state reconstruction (Puttick 2016), particularly if survival to the present day was conditional on biogeographic states themselves, given these models often exhibit poor performance in the face of character state trends through time (Finarelli and Flynn 2006; Slater et al. 2012; Raj Pant et al. 2014). As well, diversification rate estimation through the calculation of speciation and extinction rates, a common motivation for generating dated phylogenies, remains controversial in systematics (Moore et al. 2016; Meyer et al. 2018; Louca and Pennell 2020). Many investigators have pointed to the importance of fossil taxa (Upham et al. 2021) for improving inferences of diversification rates, especially the relative contributions of speciation and extinction. As in model-based biogeography, extinction likely cannot be estimated meaningfully with extant-only phylogenetic datasets (Rabosky 2010; Louca and Pennell 2021); even for speciation rate estimates, extant-only trees contain the most information at the present with decreasing power to estimate speciation dynamics closer to the root (Title and Rabosky 2019; Louca and Pennell 2020; Upham et al. 2021). Understanding the impact of fossil inclusion is therefore of broad importance to investigators, providing a means to understand the potential limits of interpretation when macroevolutionary models are built for clades of life for which direct fossil evidence is not available.

The rosid order Fagales, comprising seven extant families, 34 genera, and approximately 1,175 species distributed primarily in the temperate areas of the Northern Hemisphere (Stevens 2001 onwards), stands out for its extraordinary fossil record, perhaps the best for any large clade of flowering plants, including wood, leaves, pollen, and fruits for most of its families. The ecological dominance of Fagales in temperate forests and their economic importance have led to numerous investigations into its evolutionary relationships, ecology, morphological and anatomical variation, and the fossil record. The phylogeny of Fagales has been particularly well studied, with most studies focusing on the evolutionary relationships among its extant, currently recognized families or the specific position of certain enigmatic extant taxa (e.g., Manos & Steele 1997, Li et al. 2002, 2004, Cook & Crisp 2005, Herbert et al. 2006). Inferred relationships within Fagales over the past few decades have proven largely stable, with the major point of contention being the placement of Myricaceae, whose position is of evolutionary interest as one of three groups in Fagales to include species that engage in symbioses with nitrogen-fixing bacteria (Ardley and Sprent 2021). Phylogenies have been additionally used to understand how different fruit morphologies drove the diversification of the order (e.g., Xiang et al. 2014, Larson-Johnson 2015) and as a model to study the impact of different node calibrations (Sauquet et al. 2012).

Most phylogenetic studies of the Fagales have followed the predominant practice in divergence time estimation in plants: the fossil record is largely used as a source for calibration points for node-dating molecular-based trees. The crown age estimated for the order in these studies varies widely, but most studies place the origin of Fagales in the early (e.g., Magallón et al. 2015, Tank et al. 2015, Xiang et al. 2014, Xing et al. 2014) or late (e.g., Cook and Crisp 2005, Larson-Johnson 2015) Cretaceous. These fairly old crown ages reflect the fossil record of Fagales, as several fossils assigned to the order are dated to the earliest part of the Late Cretaceous (e.g., *Soepadmoa cupulata* [94–90 Ma], *Calathiocarpus minimus* [91.1–86.3 Ma], *Archaefagacea futabensis* [89.3–86.3 Ma]). Nevertheless, empirical tests in Fagales (Sauquet et al. 2012) demonstrate how the use of taxonomic fossil assignments, possibly overly precise analytically and therefore not adequately capturing the uncertainty in fossil placement, as well as different inference methods, greatly impact the resulting divergence ages. In that study, the use of eight calibration scenarios and two different methods resulted in a crown age range varying by a factor of 2—from ca. 120 to 67 Ma— nearly spanning the entire Cretaceous epoch (Sauquet et al. 2012).

Larson-Johnson (2016) is the only study to date to use total evidence dating approaches across Fagales. This study used 27 fossil taxa, a morphological matrix of 89 characters and five molecular loci to construct a well-supported timeline for the clade. However, despite the careful construction of the morphological matrix, several of the early fossil records for Fagales were not included, with unknown impacts on inference. Beyond this, there has been rapid progress in documenting the history of Fagales with what is now the oldest known Fagales fossil (*Soepadmoa cupulata,* Gandolfo et al. 2018) published after Larson-Johnson (2016); hence, there is a need for an up-to-date review of the fossil record and particular scrutiny of the earliest evolutionary history of Fagales. Finally, some methodological considerations in this previous study merit renewed evaluation. The morphological matrix of Larson-Johnson (2016) contained substantial missing but ultimately available morphological data for both fossil and extant taxa. That study also opted to impose a hard crown age constraint at 96.6 Ma, based on the age of the oldest fossil used, instead of using an age interval as is more typical, causing the crown age of Fagales to not be directly estimated from the data in that study. Given the sudden appearance of several distinct fossil lineages, some more closely related to extant taxa, almost immediately after 96 Ma, it seems likely that relaxing that root constraint with better early fossil sampling will improve our knowledge of origins and early diversification of the group.

A fully integrated framework for uncovering macroevolutionary processes not only includes best use of fossil data but updated and improved phylogenetic hypotheses. Previous molecular phylogenetic studies of Fagales relied almost exclusively on a few plastid loci and the nuclear ribosomal internal transcribed spacer (ITS) (e.g., Cook and Crisp 2005, Larson-Johnson 2015, Tank et al. 2015, Xiang et al. 2014, Xing et al. 2014), both high-copy loci in plants with unique evolutionary properties (Rieseberg and Soltis 1991; Alvarez and Wendel 2003) that are potentially problematic for phylogenetic inference. Our perception of the prevalence of gene tree and species tree incongruence (Maddison 1997) has only been magnified by the increasing accessibility of phylogenomic datasets (Folk et al. 2018).

Together with the well-documented prevalence of differing evolutionary histories and processes in different genomic compartments (Rieseberg et al. 1996), this has led to a shift in plant phylogenetics towards low-copy nuclear loci, sometimes leading to surprising relationships when compared to plastid phylogenies [Asteraceae: (Mandel et al. 2019), COM clade of the rosids: (Sun et al. 2015)]. Hence, particularly given the conflicting positions of Myricaceae within Fagales, a low-copy nuclear dataset is desirable to potentially decrease levels of uncertainty about critical relationships in this important plant group.

Here, we reconstruct the phylogeny and infer the biogeographic and diversification history of Fagales, building the most extensive fossil-based total-evidence analysis to date, particularly improving documentation of the earliest history of diversification of the clade compared to previous efforts. Employing more comprehensive taxon sampling, we use a new panel of low-copy nuclear loci and an expanded morphological matrix including a large fossil dataset to apply a total-evidence dating approach using the FBD model (Ronquist et al. 2012, Heath et al. 2014). Our main goals, divided across empirical and methodological aims, are to: 1) reevaluate how a nuclear dataset and an expanded morphological matrix including fossil taxa influence the inference of relationships within Fagales; 2) estimate divergence times in Fagales using a total-evidence fossilized birth-death model and compare the impact of data and assumptions with previous studies; 3) reconstruct the biogeographical history of Fagales with the inclusion of fossil taxa and compare the result with reconstruction approaches without fossils; and 4) reconstruct the history of diversification of Fagales and evaluate the impact of information from the fossil record on diversification estimation.

## Materials and Methods

### Morphological data

We reviewed the literature to assemble a comprehensive fossil sampling for Fagales, expanding the fossil taxon sampling and morphological data matrix of Larson-Johnson (2016), totaling 33 extinct fossil species (adding *Alnus clarnoensis* [age estimate 44–48 Ma], *Antiquacupula sulcata* [83.6–86.3 Ma], *Archaefagacea futabensis* [86.3–89.3 Ma], *Betula leopoldae* [47–48.6 Ma], *Endressianthus miraensis* [65.5–83.5 Ma], *Palaeocarya clarnensis* [37.8–47.8 Ma], *Paraengelhardtia eocenica* [47.8–56 Ma], *Protofagacea allonensis* [83.6–86.3 Ma], and *Soepadmoa cupulata* [90–94 Ma]) (summarized in Supplemental Material Table 1). Parsimony-informative morphological characters were identified from the literature (Jordan and Hill 1999; Oh and Manos 2008; Kooyman et al. 2014; Hermsen and Gandolfo 2016), and we assembled a morphological matrix comprising 105 characters, including new information for 16 characters from wood anatomy, leaf morphology, and reproductive morphology, such as flower and inflorescence structure, and fruit and pollen morphology, expanding on the matrix of Larson-Johnson (2015). A concerted effort was also made to fill in available missing data from extant species and correct erroneous data using literature and specimen observations (available at https://github.com/carol-siniscalchi/fagales). Eighty-eight species were scored, including the 33 fossil taxa plus at least one representative of each extant main lineage of Fagales, interpreted here as currently recognized genera. Three representative species of *Quercus* were included to account for possible non-monophyly of the genus, as previously indicated in some studies (Oh and Manos 2008; Yang et al. 2021b). A Mk model of character substitution (Lewis 2001), considering equal rates of transition among character states, was applied to this morphological matrix. We gathered data on the stratigraphic range of extinct fossil taxa and the fossil record of extant genera to provide a minimum and maximum age covering all taxa in the analysis (Supplemental Material Table 1). *Hamamelis japonica*, earlier selected by Larson-Johnson (2016) as the outgroup, was not included here for two reasons: (1) as a very distant sister to the rosid clade, *Hamamelis* makes a poor outgroup comparison to this deeply nested rosid lineage, and (2) FBD analyses usually use rooting information within the ingroup (Gavryushkina et al. 2017; May et al. 2021) rather than an outgroup because outgroup sampling strategies violate FBD assumptions pertaining to diversification priors.

### DNA sequence data

The DNA sequence data used here were extracted from the set of 100 low-copy nuclear loci used in a project (Folk et al. 2021a; Kates et al. 2021) that sampled ca. 15,000 species of the nitrogen-fixing clade of angiosperms, of which Fagales are a part. Specimens were processed using a sequence capture approach targeting loci optimized across the rosid clade, with further details available in Folk et al. (2021a). In most cases, we selected DNA data for the same species chosen for the morphological scoring; in a few cases where the same species was not available in the molecular dataset, we used the literature to select a closely related species of the same genus. The DNA sequences were assembled with aTRAM 2 (Allen et al. 2015, 2018), and, on a per-taxon basis, loci identified as potentially paralogous (i.e., presenting more than one assembled copy for one individual) were removed using the gene-tree-based pipeline of (Yang and Smith 2014). Sequences obtained for each locus were aligned with MAFFT (Katoh et al. 2009).

The use of a relatively large number of genes (e.g., the original 100 loci assembled here) is usually considered computationally intractable with FBD analyses (May et al. 2021) and previous work has likewise shown that for divergence time estimation, moderate numbers of loci are effective with diminishing returns in larger datasets (Zheng and Wiens 2015); hence, we used a “gene shopping” approach (Smith et al. 2018) to choose genes, using data completeness as the optimality criterion. Aligned gene matrices containing all sampled Fagales in the larger study by Kates et al. (2022) (up to 775 taxa per locus) were trimmed to match the 55 extant species contained in the morphological matrix, and the four loci with the highest species coverage were selected. These four loci were concatenated using the pxcat function in the package phyx (Brown et al. 2017), resulting in a matrix with 36 taxa and 5314 nucleotides. The concatenated molecular matrix was considered a single partition to simplify computation, and the GTR + Γ model was used to estimate character substitution.

### Phylogenetic analysis

The morphological and molecular datasets were combined in a Bayesian analysis in RevBayes (Höhna et al. 2014, 2017). Tree topology and divergence times were simultaneously estimated using the total-evidence FBD model (Heath et al. 2014). The age of origin of Fagales was conditioned on a uniform distribution with the minimum age defined as the maximum age of the oldest fossil assignable to Fagales (*Soepadmoa cupulata,* 94 Ma) and the maximum age defined as the maximum age of direct fossil evidence for the eudicots (125 Ma) (Anderson et al. 2005; Magallón et al. 2015). An uncorrelated molecular clock model was applied to the analysis, and no topological constraints were considered.

Ten parallel runs were conducted for 500,000 generations, with sampling at every 50 generations. Each run had a burn-in period of 5,000 generations, with a parameter tune-in every 50 generations. Convergence was estimated manually, and parameter sampling was estimated using estimated sample size (ESS) values in Tracer v1.7.1 (Rambaut et al. 2018), considering ESS = 200 as the minimum for adequate parameter sampling. A combined log of all ten runs was generated by randomly selecting 1,000 trees from each individual run log. The trees sampled in the combined log were summarized using a maximum clade credibility (MCC) approach. All RevBayes scripts and input files are available at GitHub (https://github.com/carol-siniscalchi/fagales).

To understand what factors drove divergence time differences in our analysis, we implemented two extra analyses to compare our results with Larson-Johnson’s (2016) previous Fagales phylogeny. These extra analyses included a subset of our initial data, reduced to match the taxon sampling of the earlier study, yielding 58 taxa, 34 of which are extant species, and 24 fossil taxa. Notably, Larson-Johnson (2016) did not independently estimate the age of Fagales, but instead constrained the crown age of Fagales at 96.6 Ma based on the oldest direct fossil evidence of Fagales. To test the effects of specifically constraining the crown age, we ran two analyses on this taxon subset: one where the maximum crown age was set as 96.6 Ma, as used in the previous study, and the minimum age as 91 Ma (maximum age of the oldest fossil in this dataset), and another where the maximum age was set as 125 Ma, as in the full analysis, and the minimum age as 91 Ma. One independent run was carried out for each analysis, with the same settings as above.

### Biogeographical reconstructions

We used a model-comparison approach implemented in BioGeoBEARS to reconstruct ancestral areas, including fossil tips, to directly provide novel evidence for biogeographic reconstructions not yet attempted in previous studies of Fagales. The distribution ranges of all taxa were coded based on an eight-area system (Fig. 2). Biogeographic regions were delimited largely to follow continental landmasses (Fig. 2). Eurasia was divided into a boreal + European area, a western Asian area, and an East Asian area. North America (including Central America) and South America were recognized as distinct regions, as were Africa, an Australia–New Zealand area, and a Malesia area. The treatment of New Caledonia was challenging, as this area has unique Fagales diversity, including affinities with both the northern and southern hemispheres. After trials treating it as combined with either the Malesia area or Australia–New Zealand, or as a separate region, we selected the first option because it showed the least reconstruction uncertainty (although overall results were similar despite differing uncertainty), a treatment supported particularly by shared diversity in Casuarinaceae and results from a recent regionalization analysis across vascular plants (Carta et al. 2022).

For extant taxa, the coding was based on their current distribution range as determined by consulting the Plants of the World Online database (https://powo.science.kew.org), while for fossil taxa, the collective occurrences documented for each taxon were considered as their distribution range. Three models were tested: DEC, DIVALIKE, and BAYAREALIKE, and “j” parameter models (representing founder speciation events) were excluded following the arguments of (Ree and Sanmartín 2018). The Akaike Information Criterion (AIC) was used to select the best-fitting model, and the optimal DEC model was used for further analysis.

### Mean annual temperature reconstruction

We implemented a standard ancestral reconstruction approach with mean annual temperature, the climatic variable best studied in fossil plant sites, using model comparison in the R package phytools. Mean annual temperature estimates for modern taxa were derived by downloading GBIF and iDigBio data for each species clipped to the extent of native occurrences documented in POWO and extracting BIO1 data from BioClim v1 (Hijmans et al. 2005). Mean annual temperature estimates for several of the fossils used were obtained from previous paleoclimatic reconstructions at several well-studied fossil flora localities (available at https://github.com/carol-siniscalchi/fagales). Model comparison used a standard Brownian model, an Ornstein-Uhlenbeck model, and an early-burst model; based on AIC, the Ornstein-Uhlenbeck model was favored.

### Diversification analysis

We used RevBayes to estimate diversification rates through time making full use of the fossil data, applying an episodic diversification rate model (Höhna 2015), which periodizes the diversification history in bins and estimates speciation and extinction in each bin. We followed recommendations in RevBayes documentation to set up a trial of dividing the complete timeframe of the MCC tree into either 10 intervals (∼11.3 Ma each) or 20 intervals (∼5.9 Ma each), estimating diversification rates and credibility intervals in each of these time intervals. Ultimately, we selected the 20-interval analysis because it reveals more detail with similar estimates of uncertainty. The tree and root age were set as fixed values from the main FBD analysis. Incomplete taxon sampling at the species level was modeled using taxon totals for Fagales from the Angiosperm Phylogeny Website (Stevens 2001 onwards), assuming uniform sampling within genera. Analyses were run for 10 million generations, with tuning intervals of 5,000 generations. The R package RevGadgets v.1.0.0 (Tribble et al. 2022) was used to plot the speciation, extinction, net diversification, and relative extinction rates.

### Assessing the impact of fossil taxon inclusion

Ancestral state reconstruction and reconstruction of past diversification dynamics are popular applications in contemporary systematics. While increased inclusion of fossil taxa in such analyses has been advocated to overcome the limitations of extant-only phylogenies (Stadler 2013; Rabosky 2016), studies have primarily focused on simulation (Didier et al. 2017; Mitchell et al. 2019). Empirical assessments of the benefits of using fossil data directly have been few, and not all simulation assessments of the benefits of fossil inclusion have been positive (Beaulieu and O’Meara 2022). We therefore performed a reanalysis of biogeography, ancestral temperature reconstruction, and diversification on the MCC tree excluding all lineages not extending to the present. Analyses parameters were otherwise identical to those described above, to simulate the sampling practices of extant-only trees. Our specific aims were to assess whether (1) biased extinction of particular ancestral states imposed a directional bias towards the states of extant taxa and (2) diversification estimates were biased towards recent timescales.

### Data availability

The morphological and molecular matrices, fossil and environmental data, and scripts to perform FBD and other downstream analyses used in this study are available at https://github.com/carol-siniscalchi/fagales.

## Results

### Topological features

While our MCC tree shows relationships similar to previous studies, the most notable novel feature of the Fagales topology (Fig. 1) is the presence of a fossil-only clade formed by most of the oldest Cretaceous fossils, namely *Endressianthus miraensis, Normanthus miraensis, Archaefagacea futabensis, Budvaricarpus serialis, Dahlgrenianthus suecicus, Calathiocarpus minimus, Caryanthus knoblochii, Manningia crassa*, and *Antiquocarya verruculosa*. These taxa are broadly assignable to what has been called the *Normapolles* complex (Friis et al. 2011), and this clade emerges as sister to the rest of Fagales. The other notable topological feature of our tree was that we recovered a sister relationship between Fagaceae and Nothofagaceae. While this relationship is consistent with a previous hypothesis in which *Nothofagus* was included within Fagaceae (e.g. Cronquist 1981), most recent analyses instead have found Nothofagaceae and Fagaceae successive sisters to the remainder of Fagales (Li et al. 2004; One Thousand Plant Transcriptomes Initiative 2019). The seven extant families in the order emerge as two clades with Fagaceae and Nothofagaceae reciprocal sisters in one clade, and the other five extant families placed in the other sister clade. The Cretaceous taxa *Antiquacupula sulcata* and *Protofagacea allonensis* emerge as sister to the Fagaceae, consistent with earlier inferences from morphology (Sims et al. 1988; Herendeen et al. 1995), while *Soepadmoa cupulata* is the sister taxon to Nothofagaceae.

**Figure 1.**
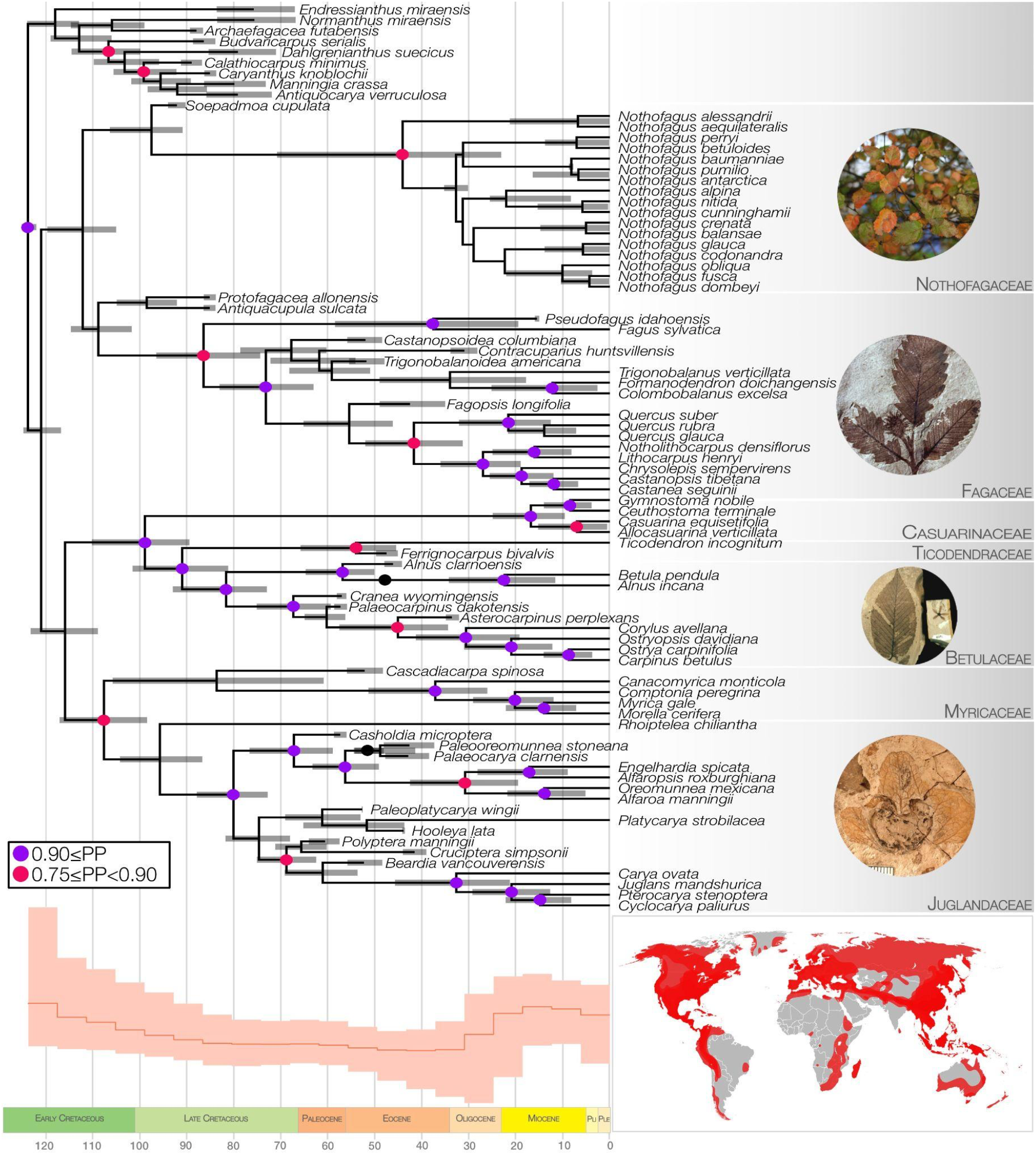
Total-evidence dating result for Fagales. Gray bars represent 95% credibility intervals; node colors represent support levels. The inset on the bottom shows net diversification through time, and the bottom right detail shows the current distribution of the order. Photo credits: Betulaceae, Fagaceae, Juglandaceae: Steven R. Manchester; Nothofagaceae: Hernan Reyes via Wikimedia.

The clade formed by the remaining five extant families is further subdivided into two clades, one formed by Casuarinaceae sister to Ticodendraceae and Betulaceae, and the other with Myricaceae and Juglandaceae sister to each other. Most of the remaining fossil taxa emerge in the tree within the families they are traditionally associated with, except for the Eocene species, *Cascadiacarpa spinosa*. The original description placed this species in Fagaceae based on fruit morphology, the only organ available. In our analyses, it emerges as sister to Myricaceae with low support (PP [posterior probability] 0.44). The FDB process allows the sampling of fossils as ancestors of extant lineages (Heath et al. 2014); in our results, *Betula leopoldae* is resolved as a sampled ancestor of the species pair *Betula pendula* and *Alnus incana* in Betulaceae, with low support (PP 0.27), and *Paraengelhardtia eocenica* is a sampled ancestor of the fossil taxa *Paleooreomunnea stoneana* and *Palaeocarya clarnensis*, in Juglandaceae, also with low support (PP 0.05).

Posterior probability values are generally low in nodes leading to fossil branches, which are dependent only on morphological evidence (Fig. 1, Supplemental Fig. 1); this is expected under the FBD and is similar to results from other recent investigations (May et al. 2021). Support values increase towards the shallower nodes, especially in clades formed by extant species, reflecting the contribution of molecular data. The crown nodes of the seven extant families have moderate (0.75 to 0.90) to high (>0.90) support values. The Fagaceae and Nothofagaceae divergence receives lower support (PP 0.7), reflecting the missing data inherent among fossil taxa with respect to these two extant families.

A notable feature of our analysis compared to all previous analyses of Fagales is the use of the FBD to produce an independent assessment of the crown age of Fagales. We obtained a point estimate of the crown age of 123.6 Ma, with the 95% highest posterior density (HPD) credible interval ranging from 121.8 to 124.9 Ma (Fig. 1, Supplemental Fig. 2). The crown ages of Fagaceae (86.3 Ma), Betulaceae (81.5 Ma), and Juglandaceae (95.7 Ma) place the respective MRCA for these families in the late Cretaceous, while Nothofagaceae (44 Ma), Ticodendraceae (53.8 Ma), and Myricaceae (37 Ma) each have their crown ages in the Eocene. Casuarinaceae has the youngest crown age, at 17.1 Ma in the Miocene. The stem ages of all extant families, however, place them in the early to late Cretaceous transition.

**Figure 2.**
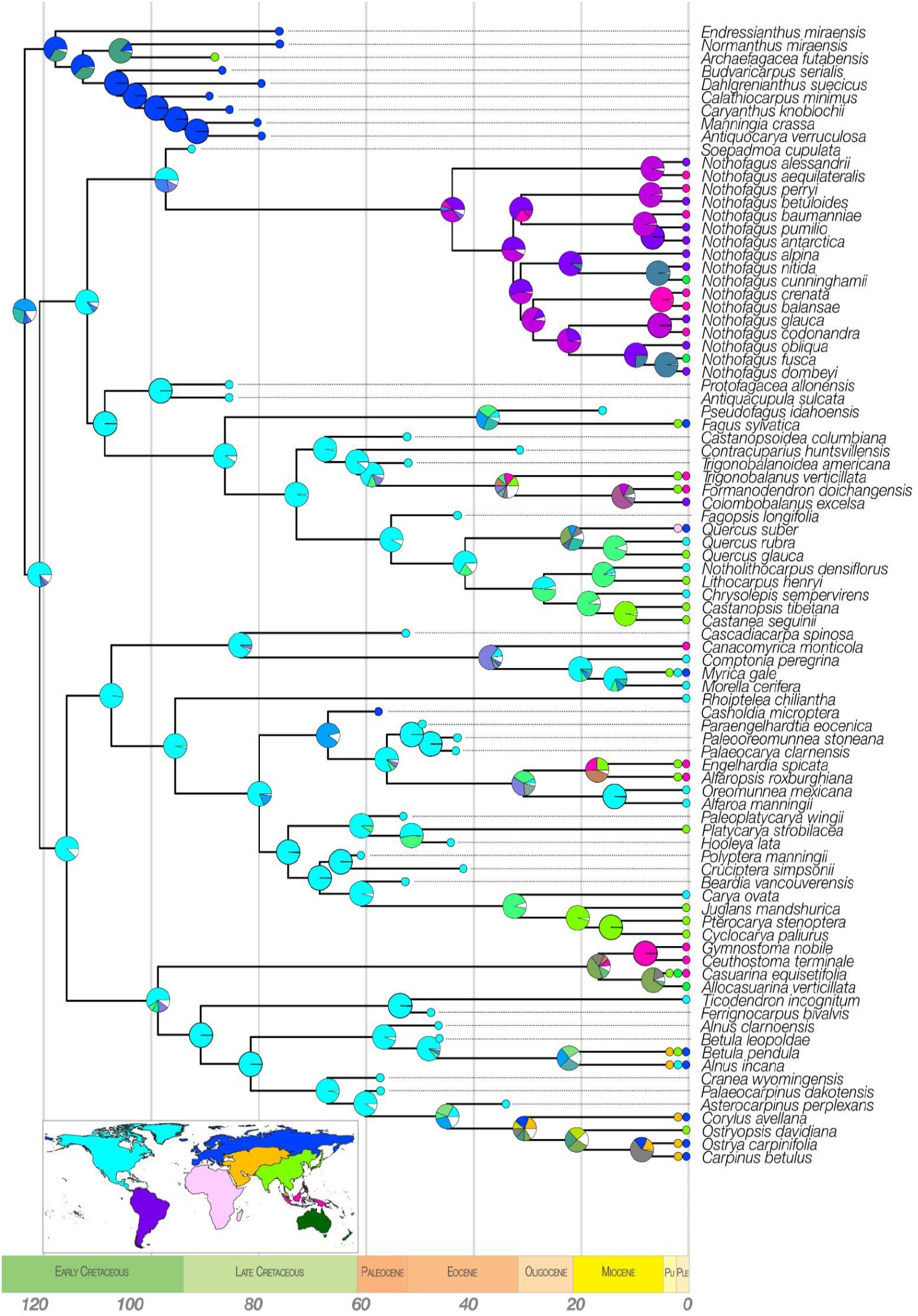
Biogeographic reconstruction under the DEC model. The inset at the bottom shows the recognized biogeographic regions with approximate extents, with colors corresponding to pie charts at nodes. Minor regions representing less than 5% of probability are not shown (white).

### Taxon sampling effects

We implemented a series of sampling and parameterization experiments to identify the cause of the older timeline we recovered for Fagales in comparison with other studies (e.g., Cook and Crisp 2005, Larson-Johnson 2016). Our tests with the reduced dataset, matching the sampling from Larson-Johnson (2016) and a narrower age interval (91–96.6 Ma) to parallel the stricter constraint used previously, resulted in an age of origin for the clade at 95.9 Ma (95% HPD 95.0–96.6 Ma, Supplemental Fig. 3), which clearly reflects the 96.6 Ma imposed maximum age; hence, the timeline of crown diversification for extant families is correspondingly much younger based on this constraint. If a wider age of origin range, following the upper bound of our full dataset, is applied to this reduced dataset, the age of origin is then estimated at 122.8 Ma (95% HPD 119.9–124.9, Supplemental Fig. 4), similar to the age found in the MCC tree of the full dataset, demonstrating that taxon sampling alone does not change the older inferred timeframe. Our full updated analysis more closely reflects typically applied constraints using the eudicot divergence (125 Ma based on earliest fossil occurrences, e.g., Magallón et al. 2015).

**Figure 3.**
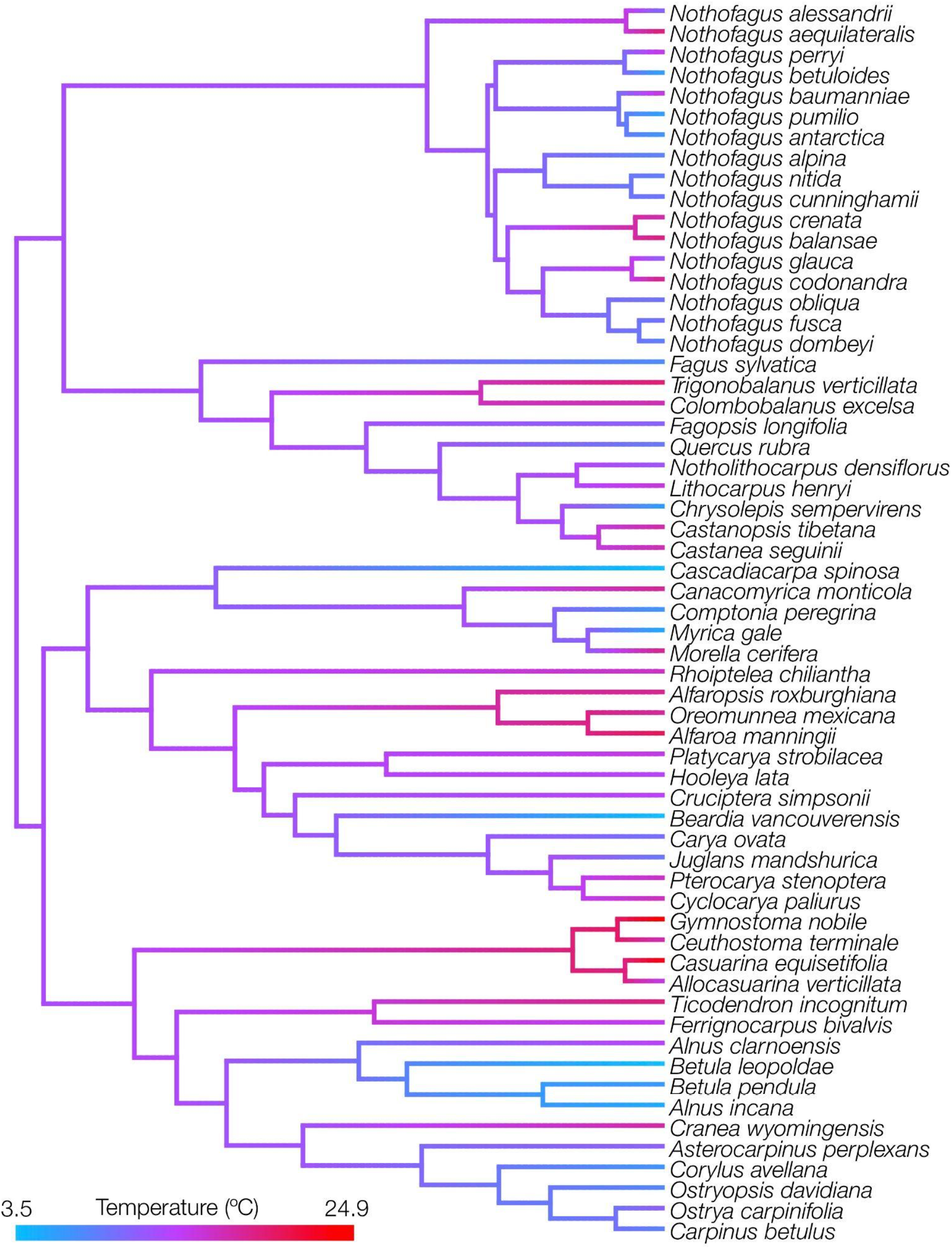
Temperature niche ancestral reconstruction; units are degrees Celsius.

### Biogeography, mean annual temperature, and diversification histories

Our biogeographical reconstruction with full taxon sampling indicates a joint area of North America and Boreal Eurasia, corresponding to a circumboreal distribution, as the probable area of origin of Fagales, with marginal probability of 43.9% (Fig. 2); a broader distribution adding East Asia to this range was next most probable at 28.7%, and no other ancestral range exceeded 5% probability. The extinct *Normapolles* clade, sister to the rest of the family, originated and largely remained in Boreal Eurasia (probability 64.5%), with only *Archaefagacea futabensis* being found in East Asia. The ancestral area of the MRCA of extant families is likely North America (probability 77.8%), indicating a range constriction. At the next split with extant representatives, both the ancestral areas of (1) the MRCA of the clade composed of Betulaceae, Casuarinaceae, Juglandaceae, Myricaceae, and Ticodendraceae, and (2) the MRCA of the clade composed of Fagaceae and Nothofagaceae are reconstructed as North America (87.8% and 85%, respectively). In general, much of the backbone was reconstructed as restricted to North America from the later Early Cretaceous until the Eocene, and this range was maintained in crown Betulaceae, Fagaceae and Juglandaceae. Nothofagaceae, unknown in the fossil record from North America, showed a possible single dispersal before 45 Ma to South America (36.2%; a second wider distribution including Malesia was next-most probable at 33.1%). The areas of MRCAs of Casuarinaceae and Myricaceae are reconstructed as combined areas (East Asia, Oceania and Malesia (42.7%), North America and Malesia (67.7%), respectively). Range expansions occurred in most families in the crown node at 30–90 Ma.

Ancestral reconstructions of mean annual temperature recovered a warm-temperate ancestor (12.8°C), a result driven largely by the richness of temperate fossil data (the extant-only analysis, discussed below, recovered a warmer ancestor). Fagales is a marginal element (and largely montane and climatically temperate) in tropical areas, and consistent with that, the reconstruction recovers colonization of tropical areas no earlier than the Eocene (Fig. 3).

Net diversification rates for Fagales showed two peaks, one a high rate at the origin of the family until ∼90 Ma, and the second increasing towards the present beginning in the early Miocene ∼25 Ma. Between these is a period of low diversification extending from 90 to 25 Ma, thus extending from the end of the Cretaceous to the Eocene, similar to the range contraction recovered in biogeographic modeling above.

### Extant taxon sampling effects

A reconstruction considering only extant taxa (Supplemental Fig. 5) shows a remarkably different biogeographical scenario. The MRCA of Fagales is reconstructed to have occurred in North America, East Asia, and South America but with very high uncertainty (probability < 13.5%). North America, the inferred origin of most lineages in the full analysis, remains here as the ancestral area of only Juglandaceae and Ticodendraceae, and the former has an earlier Cretaceous history of leaving North America than seen in the full analysis. The ancestral area of Fagaceae shifts to East Asia, and Nothofagaceae to a combined South America and Malesia area. Overall, the biogeographic pattern shifted against the rich North American fossil record and towards a scenario more reflective of modern diversity, concentrated in temperate East Asia. The timeline of dispersal out of North America is shifted earlier to a Cretaceous date, with little in common with the full analysis until the origin of extant families.

## Discussion

We present here the first full total-evidence phylogenetic analysis of Fagales, including the largest morphological dataset including fossils ever assembled for this important plant group. Our approach, combining molecular information from four nuclear markers for one representative of each extant genus, plus 105 morphological characters scored for 33 fossil and 55 extant species, recovered a largely resolved phylogeny of the extant backbone diversity of Fagales. The relationships recovered among clades recognized at the family level agree with previous studies, including monophyly of all currently recognized families. Questions remain regarding the placement of some fossil taxa, but we obtained confident placements for others, including novel results for the *Normapolles* clade. Friis et al. (2011) found their “*Normapolles* complex” assignable to core Fagales but not to any extant family. Much in accordance with our analysis, “the *Normapolles* complex probably includes mostly extinct lineages that are various kinds of stem groups among core Fagales” (Friis et al. 2011), but not in accordance with some prior interpretations, considering some of these fossil genera to be in the lineage of Juglandaceae. The ages of origin for Fagales and constituent families are generally older than in the previous total-evidence analysis of this order, but in line with many previous constraint-based analyses (e.g., Magallón et al. 2015, Tank et al. 2015, Xiang et al. 2014, Xing et al. 2014).

### Phylogenetic relationships and divergence times for extant clades

Previous phylogenetic studies of Fagales have shown mostly stable relationships among the seven constituent families. Most analyses have recovered Nothofagaceae as sister to all families, with the following families branching sequentially: Fagaceae, Juglandaceae, Myricaceae, Casuarinaceae, Ticodendraceae, and Betulaceae (Manos & Steele 1997, Cook & Crisp 2005, Sauquet et al. 2012, Xiang et al. 2014, Sun et al, 2016). The position of Myricaceae has been particularly problematic, and it is sometimes unstable with respect to Juglandaceae. Some major studies (Li et al. 2004; Magallón et al. 2015; Larson-Johnson 2016; Yang et al. 2021a) have recovered results similar to ours, with Juglandaceae and Myricaceae in a clade sister to a clade of Betulaceae, Casuarinaceae, and Ticodendraceae, with the most important remaining difference being the sequential position of Nothofagaceae and Fagaceae. These previous studies mostly shared the same set of plastid markers: *atpB, rbcL, matK, atpB-rbcL, trnL*, and *trnL-trnF*. Two studies additionally used the mitochondrial locus *matR* (Larson-Johnson 2016, Sun et al. 2016), and four used nuclear ribosomal regions (ITS: Larson-Johnson 2016, Cook & Crisp 2005, Sauquet et al. 2012; 18S/26S: Magallón et al. 2015). Given that these studies largely re-used a shared set of previously generated sequences deposited on GenBank, the similarities among topologies are not surprising.

Our study is the first to use low-copy nuclear markers to study relationships in Fagales with this level of complete genus-level sampling. The relationships we found are similar to those presented by Larson-Johnson (2016) and Magallón et al. (2015) using ITS, with one major difference: we recovered Fagaceae and Nothofagaceae sister to each other in a clade sister to the large clade of the remaining extant families. This relationship has lower support, apparently reflecting the unstable position of extinct taxa for which limited morphological information is available rather than uncertainty regarding the extant taxa.

The study of Fagales most directly comparable to ours is Larson-Johnson (2016), which is the only to include fossils as tip taxa. Using this study as a baseline, we greatly expanded the morphological matrix to include new characters and re-coded all terminals to improve data completeness and address minor coding issues. A particular goal of this renewed look was to better depict the earliest Cretaceous history of Fagales, represented by fossils omitted or not yet published and available for the earlier study (Larson-Johnson 2016). Given our concerns with this earliest history, despite rigorous data curation, Larson-Johnson (2016) constrained the crown node of Fagales at 96.6 Ma rather than letting the age of origin be informed by an age interval in tandem with the included fossils. The end result is compressed early branching within Fagales (Larson-Johnson, 2016: Fig. 3). We therefore applied the alternative approach that more fully encapsulates the principle of total-evidence dating (Ronquist et al. 2012). Fossils are normally seen as representing evidence of the *minimal* possible age of when a certain organism was alive because it is highly unlikely that the very earliest representative would happen to be in the right place and circumstances to be fossilized and subsequently recovered by a paleontologist. Under this principle, the decision of constraining the age of origin (96.6 Ma) so close to the age of the fossils (>90 Ma) is against typical practice. Because our initial results showed a much older age of origin of Fagales than that presented in Larson-Johnson (2016), we carried out a test with a reduced dataset, simulating the 96.6 Ma root constraint. Our results show that the decision to constrain the age of origin to a narrow interval is solely responsible for the timescale of the whole tree, independent of taxon sampling. This methodological decision largely constrains the rest of the Fagales timeline; for instance, the crown ages of Betulaceae, Fagaceae, and Juglandaceae in Larson-Johnson (2016) are 10 to 20 Ma younger than the dates recovered in the full analysis here, all ∼75 Ma.

Expanding the comparison to studies using other dating approaches, those using secondary calibrations have found divergent results in the age of origin of Fagales and extant families, with some consistent with our results and others closer to the younger dates of Larson-Johnson (2016). Xiang et al. (2014), using a secondary calibration approach, found an age of origin for Fagales of 105.2 Ma, and divergence times for the families similar to those in Larson-Johnson (2016). A recent total-evidence analysis of Juglandaceae (Zhang et al. 2021) found a crown age for this family of ca. 105 Ma, indeed 10 million years older than what we found here and an older timescale than a similarly recent calibration-based dating of Juglandaceae (Song et al. 2020).

Although our results overall are consistent with an older timeline for Fagales, three family ages we recovered were notable as young estimates in particularly challenging clades: Nothofagaceae, Myricaceae, and Casuarinaceae. Sauquet et al. (2012; in the favored parameterization but not others) and Xiang et al. (2014) recovered crown ages around 70 Ma for Nothofagaceae, a family that has traditionally been significant for interpreting Southern Hemisphere biogeography. Both studies used secondary calibrations at around 70 Ma for the family, based on fossil pollen, and Sauquet et al. (2012) added further secondary calibrations in a subclade within Nothofagaceae at around 30 Ma. In our analysis, four species of *Nothofagus* have their maximum ages set at 70.6 and ten species at 33.9 Ma, based on the stratigraphic record. The earlier results were on the basis of pollen evidence of Nothofagaceae from the late Cretaceous; these have been tentatively assigned to taxonomic sections and therefore used as crown constraints (Fernández et al. 2016, Swenson et al. 2000).

For Myricaceae, Larson-Johnson (2016) recovered a crown age of ca. 55 Ma, Xiang et al. (2014) recovered ca. 70 Ma, and Sauquet et al. (2012; with the true crown node of extant taxa unsampled due to not sampling *Canacomyrica*) recovered a crown age of 24.9 Ma. Our age estimates (crown age 37.1 Ma) rely on the maximum estimated ages for *Comptonia peregrina* (48.6 Ma) and *Myrica* (37.2 Ma). Xiang et al. (2014) used *Comptonia* as a calibration point at 48 Ma, while Larson-Johnson set a range between 48 and 37 Ma for the entire Myricaceae, based on fossil *Comptonia*. The unstable placement of Myricaceae in other analyses may have further contributed to this variance. We note in this context that the stem ages of all extant Fagales families place them in the early to late Cretaceous transition, meaning that stem placements of these fossils would be consistent with the foregoing older fossil evidence even where crown dates would not be.

### Biogeography and diversification

To our knowledge, this is the first thorough analysis of biogeography for Fagales as a whole. The primary focus in past work has been at the genus level, with the only major previous molecular studies at the family level being those of (Manos and Stanford 2001) for Fagaceae and (Song et al. 2020) for Juglandaceae, with equivocal results for the ancestral area of each family. We found evidence here for an ancestral circumboreal distribution for Fagalean forests occupying northern Eurasia and North America (Fig. 2), a circumscription similar to the classic Arcto-Tertiary hypothesis (Engler 1899, 1905), although earlier in date. This broad ancestral distribution was broken up soon after the crown radiation (although not simultaneously; cf. Xiang et al. (2000)) with survival only in North America in the stem of every extant lineage recognized at the family level and dispersals out of this region (as represented by extant taxa) not prevalent until the Eocene. Consistent with an early circumboreal distribution of Fagalean forests was our finding of a warm-temperate temperature niche as the ancestral state for Fagales. With an ancestral temperature similar to that of the contemporary mid-South region of the United States (12.9°C), this reconstruction suggests deep origins of the temperate niche in Fagales. Unlike the biogeographic reconstruction, ancestral temperature reconstruction proved less sensitive to the inclusion of fossil taxa, most likely because paleoclimate reconstructions at collecting localities for Fagales are largely consistent with their climate niche today.

The biogeographic scenario of broad ancestral origin, narrowed distributions, and an acceleration of dispersals in the Eocene is in agreement with our diversification estimates. We find evidence of an early burst of diversification contemporary with the origin of stem lineages to all modern families as well as several extinct lineages. A decrease in net diversification (Fig. 1) towards the end of the Cretaceous is associated with the loss of substantial Cretaceous generic diversity and range contraction. Generic diversity of Fagales begins to appear at the Paleocene-Eocene boundary but this is not associated with increased diversification. Diversification rates increase again towards the end of the Eocene or early Miocene, somewhat later than the many Eocene biogeographic dispersals of Fagales but consistent with a spread of cooler habitats during this period, resulting in largely modern distributions by the end of the Miocene. This diversification scenario is novel to our knowledge, but our finding of early elevated diversification is similar to that of Xiang et al. (2014), who found evidence for a rapid early radiation of genera of Fagales and their seed traits. A fuller exploration of the recent diversification of Fagales will require greater species-level sampling to verify this recent timeline of diversification.

### The effects of extant-only analysis

Our results show that fossil data integration into macroevolutionary models dramatically impacts inference. Fossil data made the largest impact on diversification and biogeographic reconstruction. As found previously (Rabosky 2010; Louca and Pennell 2021), extant-only chronograms tend to systematically underestimate extinction rates. While the extant-only analysis (Supplemental Fig. 6) agrees with the full analysis in recovering signs of recent diversification trends from the Miocene onwards (∼25 Ma), only the full analysis (Fig. 1; Supplemental Fig. 7) could recover a clear early burst of diversification. Notably, even with a similar shape in some respects, the scaling differed between the two analyses, with the most recent time bin being ∼2-fold greater and the oldest ∼1.5-fold less in the extant-only analysis. While an explicitly aggregated diversification-through-time estimate has not been previously performed in Fagales, the results for the extant-only approach (Bouchenak-Khelladi et al. 2015, Fig. 3) likewise clearly show the highest diversification rates in the present, suggesting the exclusion of fossils is causative and not only dataset-dependent. These results suggest optimism in attempting to infer recent speciation dynamics (Title and Rabosky 2019) but caution in attempting to interpret results inferred from near the root of a molecular phylogeny (Upham et al. 2021).

Biogeographic analysis also benefited from the high information content of fossil taxa. As mentioned above, North America was overwhelmingly the ancestral distribution for stem and crown lineages of nearly all extant families in an analysis including fossil taxa. The extant-only tree (Supplemental Fig. 5), by contrast, excludes this rich North American fossil record, and accordingly reconstructs a larger role for Asia in Fagales biogeography, particularly rewriting the early history of the largest two families, Juglandaceae and Fagaceae, both highly diverse in Asia today. In consideration of the fossil evidence, this Asian species richness is likely the result of differential extinction. An overrepresentation of extant taxa in current refugial regions may therefore be a systematic source of bias in biogeographic modeling, suggesting that fossils are particularly useful in biogeographic reconstruction in the face of extinction.

## Concluding Remarks

Our study offers an improved view of the history and biogeography of Fagales, an important angiosperm group representative of the history of today’s temperate forests. The phylogenetic relationships inferred here are well supported and largely in accord with previous molecular studies, and we were able to place Myricaceae, which have been problematic, with confidence. More broadly beyond Fagales, a stronger fossil-based framework and a series of missing data experiments demonstrated the specific benefits of fossil inclusion. Fossil exclusion, by contrast, results in relatively predictable effects on reconstructed macroevolutionary scenarios that overemphasize recent diversification and refugial regions. We can consolidate the foregoing in a series of recommendations for those using extant-only data:

1. Be aware of the limitations of ancestral state reconstruction, especially in biogeographical analyses with biased extinction across geographical areas, as expected in many empirical datasets.
2. Rather than mere random error, extant-only biogeographical reconstructions may be biased directionally towards an ancestral scenario over-representing present-day refugial areas.
3. As seen in previous simulation studies, extant-only data are better equipped to detect relatively recent bursts of diversification, but tend to reconstruct generic diversification scenarios, such as flat trends near the mean in the earliest history of a clade, imposing clear limits on interpretation.
4. Fossil data offer a path to ameliorate limitations in the interpretation of diversification analysis when either early clade history or estimating extinction itself is the objective.

## Supplementary Material

All data files, scripts and additional figures are available at https://github.com/carol-siniscalchi/fagales.

## Funding

This work was supported by the National Science Foundation grant number NSF DEB-1916632.

## Acknowledgments

We would like to thank Tracy Heath for help and advice regarding FDB models; C.M.S. also would like to thank the instructors of the Summer 2020 RevBayes Workshop for initial help and discussion, and Patrick Herendeen for providing photographs of Fagales fossils that were ultimately not included in the paper.

## Notes

https://github.com/carol-siniscalchi/fagales

### Competing Interest Statement

The authors have declared no competing interest.

