## Supplemental Figures for "Fagalean phylogeny in a nutshell: Chronicling the diversification history of Fagales"

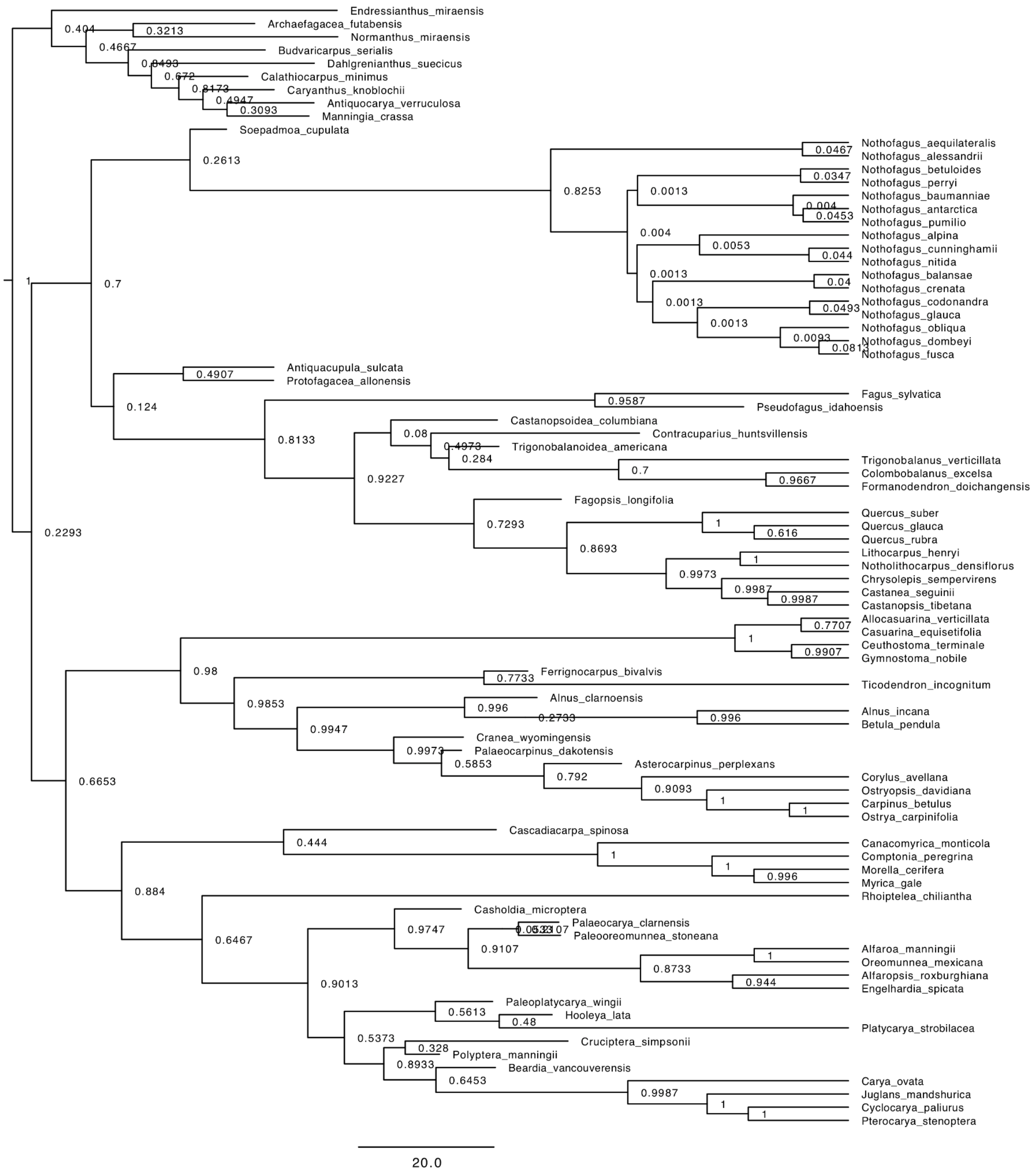


**Supplemental Figure 1.** Full tree with support values (posterior probabilities).

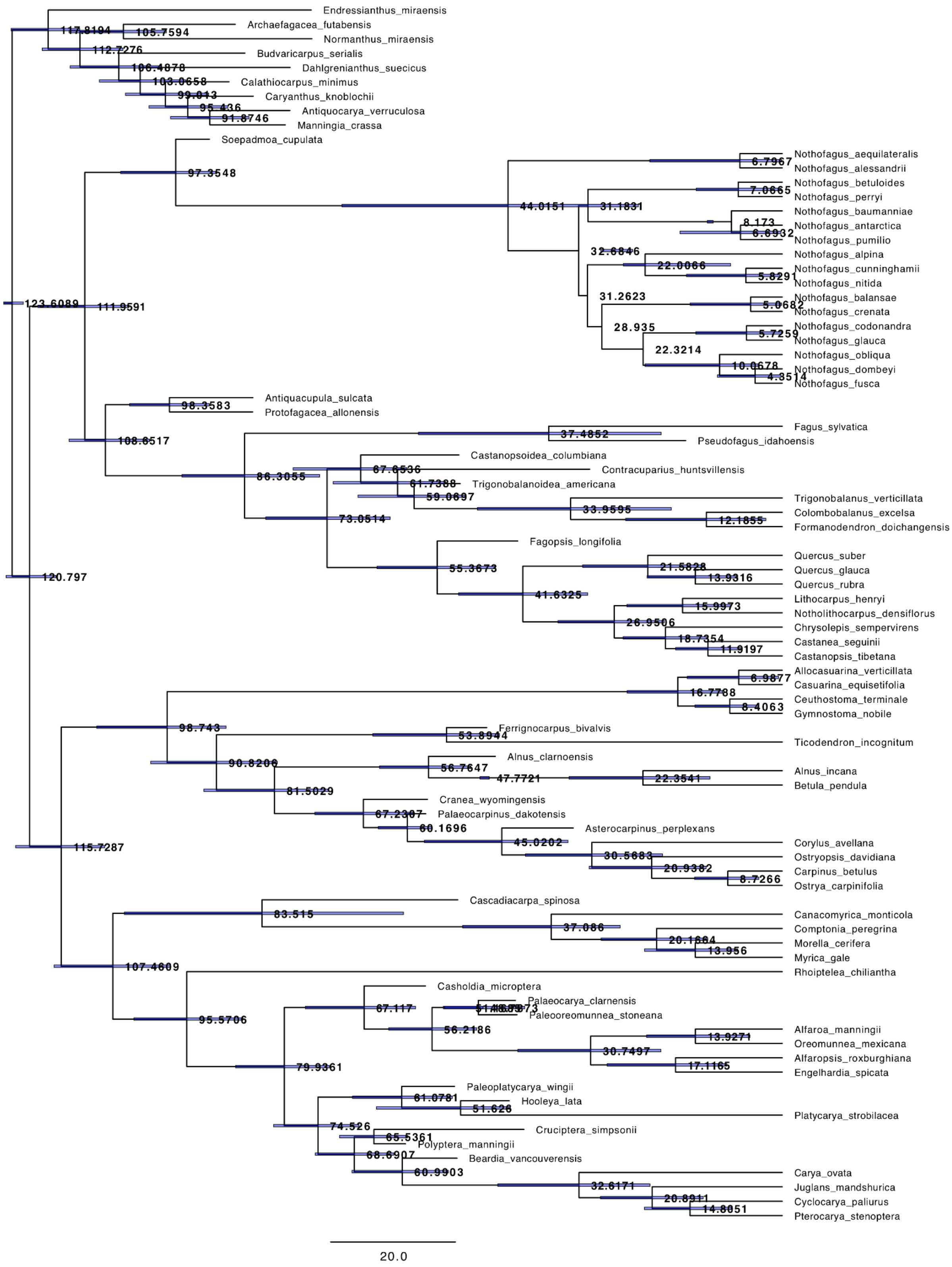


**Supplemental Figure 2.** Full tree with node ages and 95% HPD intervals.

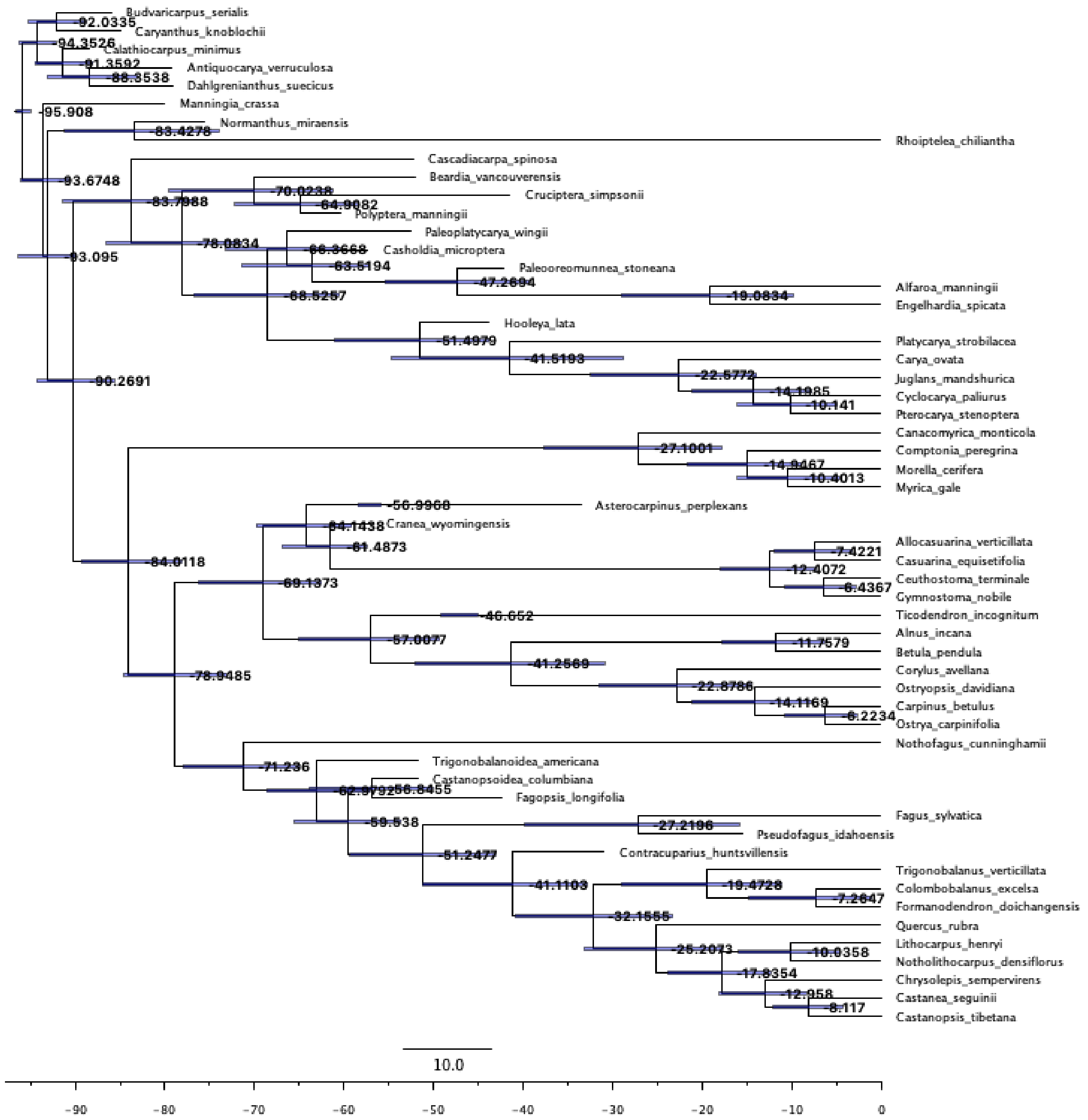


**Supplemental Figure 3.** Reduced dataset tree, matching the sampling of Larson-Johnson (2016) and using a root age interval of 91 – 96.6 Ma. Node labels represent ages and 95% HPD intervals.

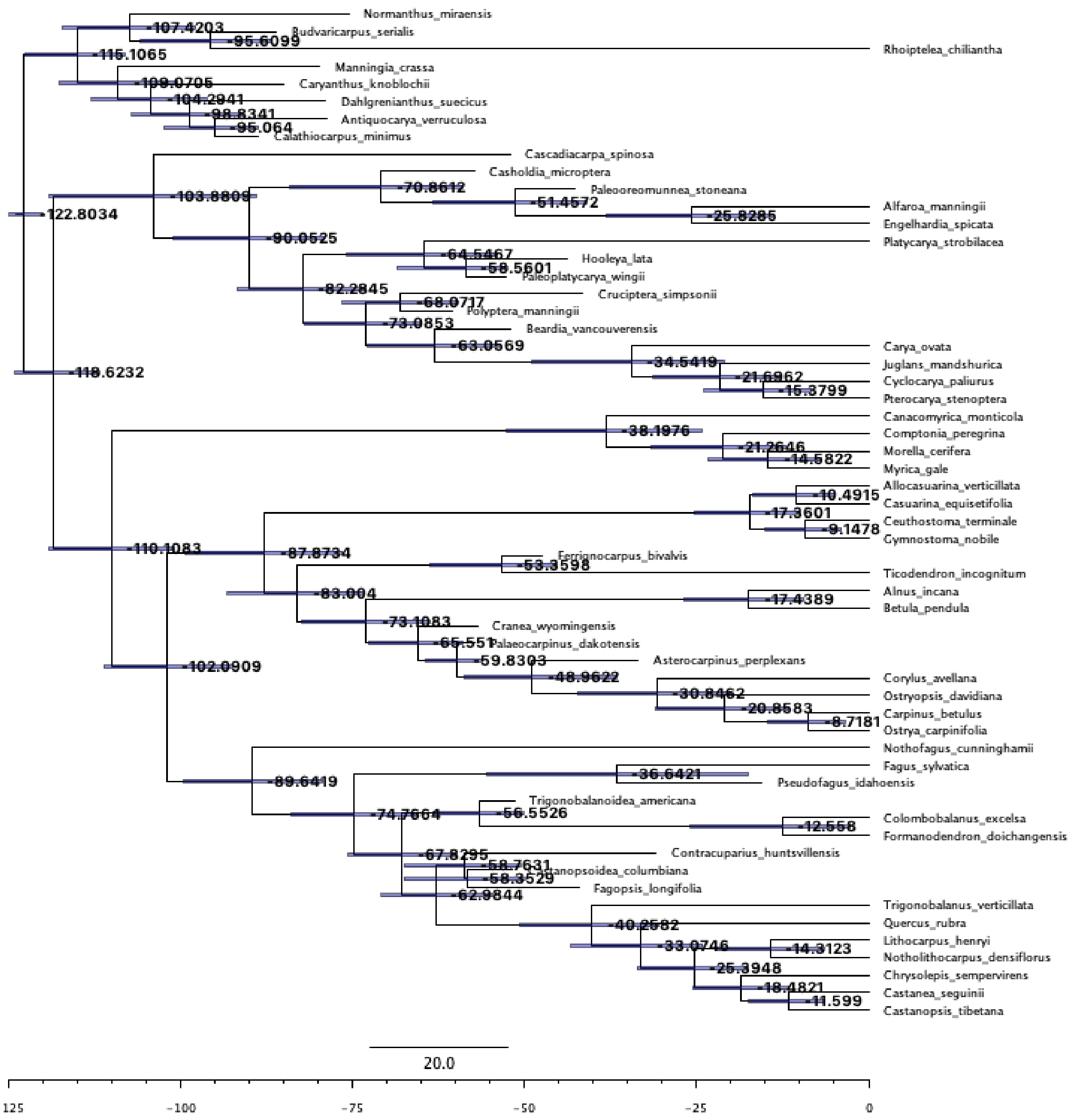


**Supplemental Figure 4.** Reduced dataset tree, matching the sampling of Larson-Johnson (2016) and using a root age interval of 91 – 125 Ma. Node labels represent ages and 95% HPD intervals.

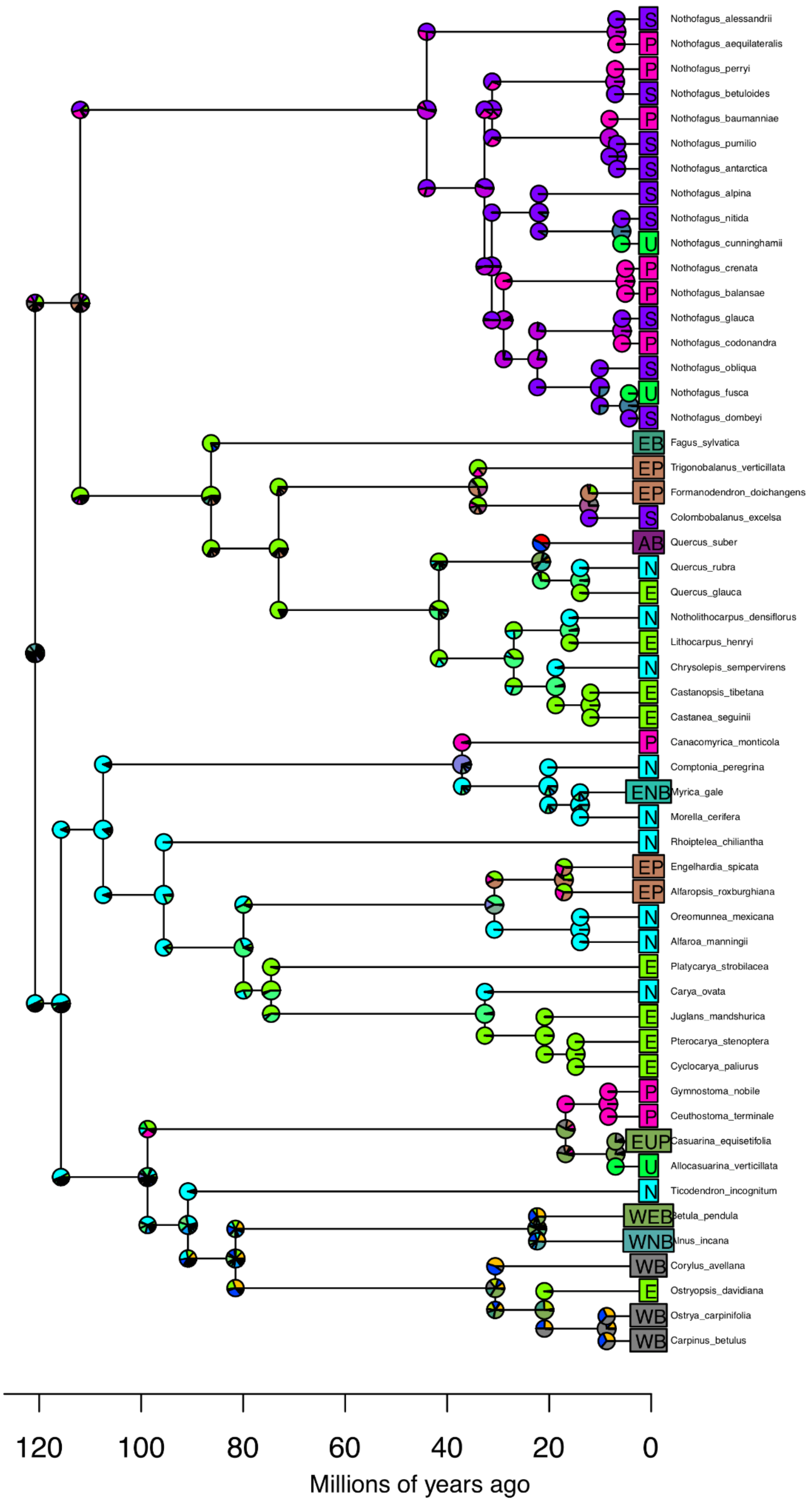


**Supplemental Figure 5.** Extant-only BioGeoBEARS run. Color scheme matches the map shown in Figure 2.

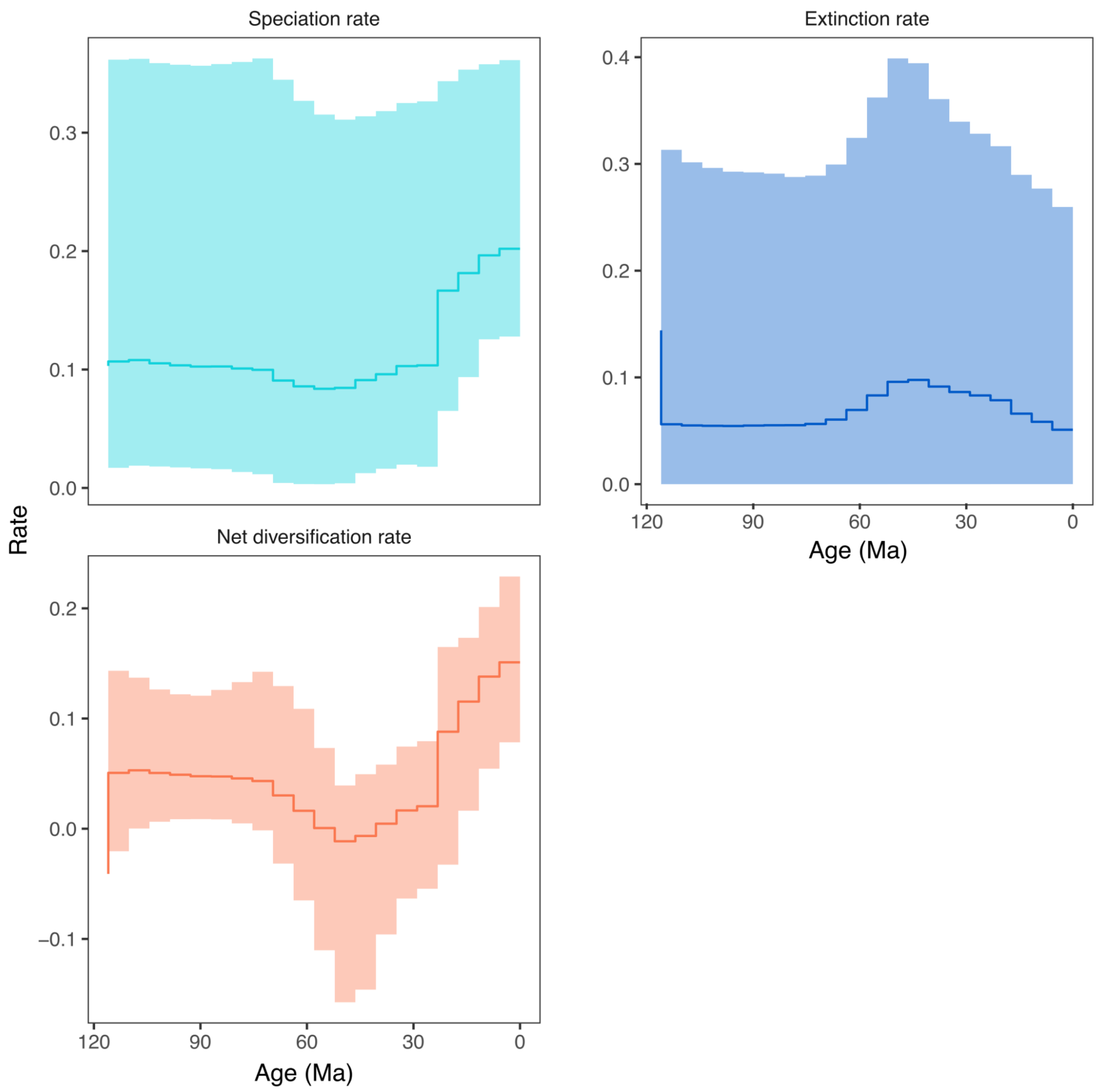


**Supplemental Figure 6.** Speciation, extinction, and net diversification plots for only extant taxa. Although some features are similar, note particularly the flat early diversification history and the change in the y-axis scale.

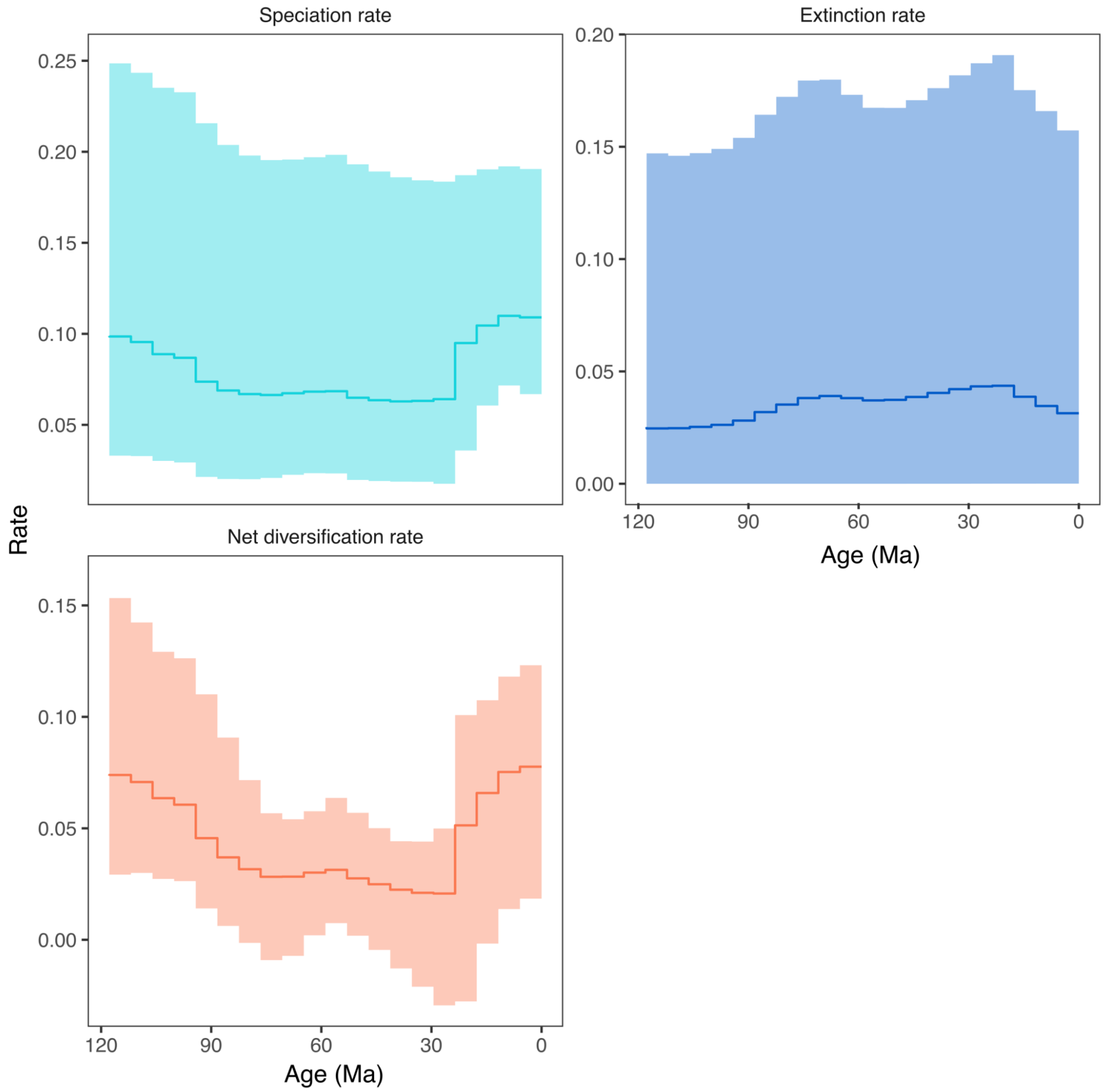


**Supplemental Figure 7.** Speciation, extinction, and net diversification plots for the full dataset. The net diversification plot is identical to Figure 1 but shown here for clarity.
