## Supplemental Table 1 for "Fagalean phylogeny in a nutshell: Chronicling the diversification history of Fagales"

| **Supplemental Table 1:** Maximum and minimum age of all taxa included in the morphological matrix, including locality of the fossil taxa used in the analysis, and mean temperature used in the reconstruction shown in Figure 3. | | | | | |
|---|---|---|---|---|---|
| **Taxon** | **Minimum age** | **Maximum age** | **Main paleolocality** | **Other Paleolocalities** | **Temperature (ºC)** |
| Alfaroa manningii | 0 | 0 | NA | NA | 21.08040 |
| Alfaropsis roxburghiana | 0 | 46 | NA | NA | 18.75210 |
| Allocasuarina verticillata | 0 | 0 | NA | NA | 14.89890 |
| Alnus clarnoensis | 44 | 48 | Central Oregon, USA | China, Europe | 14.00000 |
| Alnus incana | 0 | 48 | NA | NA | 5.18624 |
| Antiquacupula sulcata | 83.6 | 86.3 | Georgia, USA | NA | NA |
| Antiquocarya verruculosa | 70.6 | 85.8 | Scania, Sweden | NA | NA |
| Archaefagacea futabensis | 86.3 | 89.3 | Honshu, Japan | NA | NA |
| Asterocarpinus perplexans | 32 | 35 | Oregon, Colorado, Montana, USA | NA | 11.30000 |
| Beardia vancouverensis | 47.8 | 55.8 | British Columbia, Canada | NA | 3.50000 |
| Betula leopoldae | 47 | 48.6 | British Columbia, Canada | NA | 3.50000 |
| Betula pendula | 0 | 48.6 | NA | NA | 6.58120 |
| Budvaricarpus serialis | 83.5 | 88.6 | Czech Republic | NA | NA |
| Calathiocarpus minimus | 86.3 | 91.1 | Germany | Europe | NA |
| Canacomyrica monticola | 0 | 0 | NA | NA | 18.52060 |
| Carpinus betulus | 0 | 50 | NA | NA | 8.65535 |
| Carya ovata | 0 | 37.2 | NA | NA | 10.02540 |
| Caryanthus knoblochii | 83.6 | 86.3 | Scania, Sweden | Czech Republic, Georgia (USA | NA |
| Cascadiacarpa spinosa | 47.8 | 56 | British Columbia, Canada | NA | 3.50000 |
| Casholdia microptera | 55.8 | 58.7 | England | NA | NA |
| Castanea seguinii | 0 | 48.6 | NA | NA | 17.06220 |
| Castanopsis tibetana | 0 | 44.1 | NA | NA | 17.72140 |
| Castanopsoidea columbiana | 47.8 | 55.8 | Tennessee, USA | NA | NA |
| Casuarina equisetifolia | 0 | 0 | NA | NA | 24.23590 |

| | | | | | |
|---|---|---|---|---|---|
| Ceuthostoma terminale | 0 | 0 | NA | NA | 17.38570 |
| Chrysolepis sempervirens | 0 | 0 | NA | NA | 6.29411 |
| Colombobalanus excelsa | 0 | 0 | NA | NA | 17.14810 |
| Comptonia peregrina | 0 | 48.6 | NA | NA | 6.94444 |
| Contracuparius huntsvillensis | 27.8 | 33.9 | Texas, USA | NA | NA |
| Corylus avellana | 0 | 50 | NA | NA | 7.09862 |
| Cranea wyomingensis | 55.8 | 58 | Wyoming, USA | NA | 17.90000 |
| Cruciptera simpsonii | 38.8 | 44.1 | Oregon, USA | Germany, England | 14.00000 |
| Cyclocarya paliurus | 0 | 61.6 | NA | NA | 16.77820 |
| Dahlgrenianthus suecicus | 70.6 | 85.8 | Scania, Sweden | NA | NA |
| Endressianthus miraensis | 65.5 | 83.5 | Scania, Sweden | NA | NA |
| Engelhardia spicata | 0 | 0 | NA | NA | NA |
| Fagopsis longifolia | 33.9 | 49 | Colorado, USA | Montana, Washington (USA) | 11.30000 |
| Fagus sylvatica | 0 | 48.6 | NA | NA | 7.86305 |
| Ferrignocarpus bivalvis | 45 | 50 | Oregon, USA | England | 14.73000 |
| Formanodendron doichangensis | 0 | 0 | NA | NA | NA |
| Gymnostoma nobile | 0 | 51.9 | NA | NA | 24.88180 |
| Hooleya lata | 43.5 | 44.1 | Oregon, USA | | 14.00000 |
| Juglans mandshurica | 0 | 44 | NA | NA | 9.45147 |
| Lithocarpus henryi | 0 | 40 | NA | NA | 15.58500 |
| Manningia crassa | 72.1 | 86.3 | Scania, Sweden | | NA |
| Morella cerifera | 0 | 0 | NA | NA | 18.89850 |
| Myrica gale | 0 | 37.2 | NA | NA | 5.40265 |
| Normanthus miraensis | 65.5 | 83.5 | Portugal | NA | NA |
| Nothofagus aequilateralis | 0 | 70.6 | New Jersey, USA | NA | 20.02960 |
| Nothofagus alessandrii | 0 | 70.6 | Oregon, USA | Europe, Asia | 10.83000 |
| Nothofagus alpina | 0 | 33.9 | Tennessee, Kentuky, USA | NA | 8.59020 |

| | | | | | |
|---|---|---|---|---|---|
| Nothofagus antarctica | 0 | 33.9 | Wyoming, USA | NA | 6.95267 |
| Nothofagus balansae | 0 | 70.6 | Tennessee, USA | NA | 19.41150 |
| Nothofagus baumanniae | 0 | 70.6 | Colorado, Wyoming, Montana, USA | NA | 15.73330 |
| Nothofagus betuloides | 0 | 33.9 | Georgia, USA | NA | 5.87206 |
| Nothofagus codonandra | 0 | 70.6 | Idaho, USA | NA | 18.99060 |
| Nothofagus crenata | 0 | 0 | NA | NA | 17.48890 |
| Nothofagus cunninghamii | 0 | 1.8 | North Dakota, Wyoming, USA | North Hemisphere | 9.67186 |
| Nothofagus dombeyi | 0 | 33.9 | Tennessee, USA | NA | 8.76047 |
| Nothofagus fusca | 0 | 33.9 | NA | NA | 9.91588 |
| Nothofagus glauca | 0 | 33.9 | NA | NA | 11.63000 |
| Nothofagus nitida | 0 | 33.9 | NA | NA | 8.98421 |
| Nothofagus obliqua | 0 | 33.9 | NA | NA | 9.78023 |
| Nothofagus perryi | 0 | 33.9 | NA | NA | 15.24620 |
| Nothofagus pumilio | 0 | 33.9 | NA | NA | 5.87239 |
| Notholithocarpus densiflorus | 0 | 0 | NA | NA | 12.42770 |
| Oreomunnea mexicana | 0 | 19 | NA | NA | 18.41180 |
| Ostrya carpinifolia | 0 | 33 | NA | NA | 11.44090 |
| Ostryopsis davidiana | 0 | 0 | NA | NA | 8.24648 |
| Palaeocarpinus dakotensis | 55.8 | 58.7 | North Dakota, Wyoming, USA | North Hemisphere | NA |
| Palaeocarya clarnensis | 37.8 | 47.8 | Oregon, USA | Europe, Asia | NA |
| Paleooreomunnea stoneana | 37.2 | 48.6 | Tennessee, Kentuky, USA | NA | NA |
| Paleoplatycarya wingii | 52.5 | 52.7 | Wyoming, USA | NA | NA |
| Paraengelhardtia eocenica | 47.8 | 56 | Tennessee, USA | NA | NA |
| Platycarya strobilacea | 0 | 55.8 | NA | NA | 15.13250 |
| Polyptera manningii | 57 | 64 | Colorado, Wyoming, Montana, USA | | NA |
| Protofagacea allonensis | 83.6 | 86.3 | Georgia, USA | NA | NA |
| Pseudofagus idahoensis | 15 | 16 | Idaho, USA | NA | NA |

| | | | | | |
|---|---|---|---|---|---|
| Pterocarya stenoptera | 0 | 33.9 | NA | NA | 16.42620 |
| Quercus glauca | 0 | 44 | NA | NA | NA |
| Quercus rubra | 0 | 44 | NA | NA | 9.34473 |
| Quercus suber | 0 | 44 | NA | NA | NA |
| Rhoiptelea chiliantha | 0 | 0 | NA | NA | 16.46000 |
| Soepadmoa cupulata | 90 | 94 | New Jersey, USA | NA | NA |
| Ticodendron incognitum | 0 | 0 | NA | NA | 19.77950 |
| Trigonobalanoidea americana | 47.8 | 55.8 | Tennessee, USA | NA | NA |
| Trigonobalanus verticillata | 0 | 37.2 | NA | NA | 20.12310 |
